# Predicting late-onset Alzheimer’s disease from genomic data using deep neural networks

**DOI:** 10.1101/629402

**Authors:** Javier de Velasco Oriol, Edgar E. Vallejo, Karol Estrada, The Alzheimer’s Disease Neuroimaging Initiative

## Abstract

Alzheimer’s disease (AD) is the leading form of dementia. Over 25 million cases have been estimated worldwide and this number is predicted to increase two-fold every 20 years. Even though there is a variety of clinical markers available for the diagnosis of AD, the accurate and timely diagnosis of this disease remains elusive. Recently, over a dozen of genetic variants predisposing to the disease have been identified by genome-wide association studies. However, these genetic variants only explain a small fraction of the estimated genetic component of the disease. Therefore, useful predictions of AD from genetic data could not rely on these markers exclusively as they are not sufficiently informative predictors. In this study, we propose the use of deep neural networks for the prediction of late-onset Alzheimer’s disease from a large number of genetic variants. Experimental results indicate that the proposed model holds promise to produce useful predictions for clinical diagnosis of AD.

## 1 Introduction

Alzheimer’s disease (AD) is a degenerative brain disease characterized by the loss of cognitive abilities such as memory, reasoning, language and behavior. In most cases, AD ultimately leads to death.

Late-onset Alzheimer’s disease (LOAD) is the most common form of dementia (60% – 80% cases). It occurs more often in people age 60 and older. There is no known ultimate cause of LOAD. In addition, there is no effective therapeutic treatment for the disease. However, it is likely that LOAD is a complex disease which etiology is driven by both environmental and genetic components [3].

There is no single effective clinical test for LOAD. Currently, a confirmatory diagnosis of the disease is exclusively available from pathological postmortem examinations [24]. However, there is a collection of tests that are considered useful predictors for the clinical diagnosis of LOAD, such as MRI and PET images, cognitive tests, cerebrospinal and blood biomarkers, and genetic markers, among others [17].

Unfortunately, the majority of these clinical markers are strongly correlated with the progression of this disease, meaning that they would typically be more informative at later stages of the disease or are very expensive to perform on large population screens. We need better clinical tests that are capable to provide accurate predictions for the early diagnosis of LOAD. In effect, we expect experimental therapeutic and palliative interventions to be more effective at earlier stages of the disease [23].

A promising alternative for the prediction of LOAD is through genetic testing. For example, specific alleles of Apolipoprotein E (APOE) have been implicated as the largest genetic risk factors for LOAD. However, the use of the APOE4 in clinical practice has been controversial. For example, even though the odds ratio of this genetic marker has been estimated at over 3, in practice, only 1 of 4 patients with this allele progresses to the disease.

Recent advances on genome technologies have enabled the identification of several genetic variants that are associated with complex diseases [19] [18]. However, the complete understanding of the genetic architecture of most complex diseases has remained elusive [22]. Advances in this area hold the potential to contribute to the identification of novel drug targets for LOAD [11] [12].

The genetic component of LOAD has been estimated to be 79%. However, recent studies on the heritability of LOAD have estimated that common genetic variants identified by genome-wide association studies (GWAS) are only capable to explain 33% of the phenotypic variance, meaning that over 40% of the genetic component remains unexplained [21].

Recent studies have been postulated a collection of theories that should be capable to explain the missing heritability of complex diseases [19] [6]. These theories include: (1) a more comprehensive collection of genes with low effect sizes associated with the disease; the existence of genegene interactions –epistatic effects; and gene-environmental interactions, among others.

In this research, we propose to explore the hypothesis on the existence of multiple genes with low effect sizes contributing to the risk of developing LOAD. To test this hypothesis we propose to conduct computational experiments on the construction of deep learning predictive models using large collections of genetic markers that are capable to predict LOAD from this data.

Deep learning (DL) are machine learning models that are becoming increasingly popular in solving a variety of problems in medicine [7]. In recent years, DL models have shown excellent results on the examination of clinical images, approaching human level performance [10]. In this work, we propose to explore the use of dense deep neural networks (DDNN) models for predicting complex disease from genetic data.

We conducted a series of experiments on the use of deep neural networks for predicting late-onset Alzheimer’s disease using whole genome data from the Alzheimer’s Disease Neuroimaging Initiative (ADNI) project. Experimental results indicate that classification performance of ∼65% of area under the ROC curve (AUC) can be achieved with the proposed model. Further, the experiments reported here suggest that an increasing number of genetic variants hold the potential to contribute to improve the predictive capabilities of the proposed model providing that sufficiently large datasets are available.

## 2 Materials and Methods

### 2.1 Data set

Data used in the preparation of this article were obtained from the Alzheimers Disease NeuroimagingInitiative (ADNI) database (adni.loni.usc.edu). The ADNI was launched in 2003 as a public-private partnership, led by Principal Investigator Michael W. Weiner, MD. The primary goal of ADNI has been to test whether serial magnetic resonance imaging (MRI), positron emission tomography (PET), other biological markers, and clinical and neuropsychological assessment can be combined to measure the progression of mild cognitive impairment (MCI) and early Alzheimers disease (AD).

The ADNI dataset individuals have multiple possible diagnosis: Cognitively Normal (CN), Early and Late Mild Cognitive Impairment (eMCI, lMCI) and Alzheimer’s Disease (AD). The ADNI Study is divided in three different studies so far: ADNI 1, ADNI GO and ADNI 2. The ADNI 1 study consisted of 200 CN, 400 MCI and 200 AD individuals. ADNI GO extended this with 200 eMCI individuals and 500 rollovers from ADNI 1. Finally, ADNI 2 integrated those rollovers with 150 CN, 150 eMCI, 150 lMCI, and 200 AD additional individuals.

In our case the subset of data to be utilized are the Whole-Genome Sequence (WGS) samples for 812 individuals available on the ADNI database. These WGS were sampled on an Illumina Omni 2.5M chipset and containing 2,379,855 Single Nucleotide Polymorphisms(SNPs). Due to the varying nature of MCI and the uncertainty of whether the patient will progress to Alzheimer’s Disease the classification is strictly binary and as such the only samples taken for the binary classification are those that have a CN or AD diagnosis.

Additionally, the results from the International Genomics of Alzheimer’s Project [15] were also used to do feature selection, guide the learning process of the algorithm and to obtain the value for augmenting the data set artificially. The International Genomics of Alzheimer’s Project (IGAP) is a large two-stage study based upon genome-wide association studies (GWAS) on individuals of European ancestry. In stage 1, IGAP used genotyped and imputed data on 7,055,881 single nucleotide polymorphisms (SNPs) to meta-analyse four previously-published GWAS datasets consisting of 17,008 Alzheimer’s disease cases and 37,154 controls (The European Alzheimer’s disease Initiative EADI the Alzheimer Disease Genetics Consortium ADGC The Cohorts for Heart and Aging Research in Genomic Epidemiology consortium CHARGE The Genetic and Environmental Risk in AD consortium GERAD). In stage 2, 11,632 SNPs were genotyped and tested for association in an independent set of 8,572 Alzheimer’s disease cases and 11,312 controls. Finally, a meta-analysis was performed combining results from stages 1 & 2.

Most of the individuals with WGS that form part of the ADNI 1 and ADNI GO studies were also included as part of the IGAP GWAS meta-analysis, while the new individuals in the ADNI 2 study are completely independent of the IGAP. As such some experiments make the distinction between those two groups in the ADNI dataset.

### 2.2 Tools and Software

The software used to read the Variant Call Format data of the WGS and convert it to the more compact format of Binary Pedigree Files (BED) PLINK [20] [25] was used, as well as for the quality control pipeline. The code was implemented in Python 3.5, using Tensorflow [1] for the GPU backend and Keras [4] for the deep learning framework, to access the Binary Pedigree Files from python the library PyPlink [16] was used.

### 2.3 Quality Control Pipeline

When handling genetic data specific care must be taken to pre-process it by using different Quality Control methodologies, as there are some intrinsic factors in genetics that can cause methodological errors or inconsistencies between results which do not normally factor in other types of datasets. These factors can be related to the Samples, to the Markers or to the Batch effects.The pipeline described by Turner et al [26] describes some of the most common characteristics that need to be analyzed and filtered:

- Chromosomal anomalies
- Sex anomalies
- Related Samples
- Population Stratification
- Sample Call Rates
- Marker Call Rates
- Minor Allele frequency (MAF)
- HapMap Concordance
- Hardy-Weinberg Equilibrium
- Linkage Disequilibrium
- Plate measurement effects

We followed this pipeline with some adjustments. We did not perform the first two analysis as we are not concerned with the sex of the individuals and thus we are discarding the 23th chromosome. The next step was doing the sample analysis. We performed some pre-quality controls on the data (marker call rate, sample call rate and MAF) and then performing Identity-By-Descent calculations to identify those individuals that were family with an IBD sharing of more than 0.25. 8 different individuals were found to be related, and thus were added to the list to be removed. Doing a quick analysis of the IDs and the ADNI reference information we found there were no individuals from populations different from “White” in the 808 samples for the WGS, this was confirmed using Principal Component Analysis and finding no severe outliers. Due to the nature of the binary classification problem at the pre-processing stage we removed all the individuals which were assigned an EMCI, LMCI or SMC diagnosis. Considering these 3 sample filters the dataset was reduced from 808 samples to 471.

Once reduced in samples the next step was to do the call rate and MAF filtering. We begin with a dataset consisting of 471 samples with 42,908,833 variants. We performed the Sample call rate filtering with the default value of 90, which revealed that no sample were to be removed. Afterwards the Marker call rate filtering was done using a value of 99, thus removing all snps with a lower call rate than 99 and obtaining 38,517,541 markers remaining. Then the MAF was calculated and all snps with a MAF of !0.01 were also removed. 8,968,581 markers were left after the MAF thresholding.

The next step was to perform the Hardy-Weinberg Equilibrium test using a significance value of 0.05 to remove all markers with a higher value, obtaining 8,498,435 remaining markers. The last step is to perform an LD-based clumping on the data set before the pruning but without the correlated individuals. The IGAP results are then used as the association study from which to obtain the p-values for the LD-based Clump,which is then run with a p-value of 0.001 and *r*^2^ of 0.05 to obtain a list of the 1,884 best index SNP candidates which is the one that will be utilized to guide the learning procedure. The deep learning algorithms will then analyze subsets of the most significant SNPs, thus performing a more strict significance filtering later on. The HapMap concordance and the plate measurement effects were not taken into account.The first as we wanted to maximize the markers obtained from the IGAP study within the clump and the second primarily because the ADNI study already incorporates quality controls within the device procedure.

### 2.4 Neural Network Architecture

We present the general architecture of the Neural Network as well as the chosen parameters used in this research: The activation function for the neuron layers proposed is the Rectified Linear Unit “ReLU” as the gradient will be efficiently propagated and the activation of units will be small, splitting decision making across the network. Weight Dropout of 30% per Dense layer is also used to avoid the vanishing gradient problem and to avoid over-fitting. Additionally, each Dense layer is initialized using a He Normal initialization and regularized using L2 with a factor of 0.000001 Regarding the model optimization, Cross-Entropy Loss is our chosen function for minimizing the error on the training data, and the Adam default model is used as the optimizer. Additional Normalization as well as Gaussian Noise layers are also added to generalize the model after the first two layers.

Our Neural Network architecture is designed as follows:

- Input: SNPs obtained from QC Pipeline
- Dense with neurons equal to the number of inputs SNPs*, ReLU as activation, L2 Regularization and He Initialization
- Batch Normalization, Dropout Layer with 30% of inputs to drop,Gaussian Noise with 0.3 as Standard Deviation
- Dense Layer with 1024 outputs, ReLU as activation, L2 Regularization and He Initialization
- Batch Normalization, Dropout Layer with 30% of inputs to drop
- Dense Layer with 512 outputs, ReLU as activation, L2 Regularization and He Initialization
- Batch Normalization, Dropout Layer with 30% of inputs to drop
- Dense Layer with 256 outputs, ReLU as activation, L2 Regularization and He Initialization
- Batch Normalization, Dropout Layer with 30% of inputs to drop
- Dense Layer with 64 outputs, ReLU as activation, L2 Regularization and He Initialization
- Dense with 2 outputs, sigmoid activation
- Output: Prediction probability for Alzheimer’s Disease

The Convolutional network structure has two different variants, where both use 1-dimensional convolutional filters with size 5, and uses the Same padding technique to handle edges. One version uses Dropout while a different version utilizes Batch Normalization:

- Input: SNPs obtained from QC Pipeline
- 1-Dimensional Convolution with neurons equal to the number of inputs SNPs*, ReLU as activation
- Dropout Layer with 20% of inputs to drop or Batch Normalization
- 1-Dimensional Convolution with neurons equal to the number of inputs SNPs*, ReLU as activation
- Dropout Layer with 20% of inputs to drop or Batch Normalization
- 1-Dimensional Convolution with neurons equal to the number of inputs SNPs*, ReLU as activation
- Dropout Layer with 20% of inputs to drop or Batch Normalization
- 1-Dimensional Convolution with neurons equal to the number of inputs SNPs*, ReLU as activation
- Dropout Layer with 20% of inputs to drop or Batch Normalization
- Global Max Pooling
- Dense with 2 output, sigmoid activation
- Output: Prediction probability for Alzheimer’s Disease

Finally, a Support Vector Classifier using a Radial Basis Function Kernel and a Random Forest with are used to contrast and compare the different neural networks.

## 3 Simulation and Data augmentation

One of the traditional requirements for Neural networks to work is the use of a large amount of samples. Thus we wanted to consider what would happen if we had a given complex genetic disease and how the size of the data set used for training could affect the final results as the original ADNI data set is small. We began by using the PLINK simulator to generate a disease roughly distributed in a complex manner, with 912,053 SNPs, out of which there were 12,053 genes related to the disease with different Odds Ratio (A very highly-correlated SNP, some few SNPs with high-correlation, and many with low correlation). We generated 200,000 samples and then took 100,000 independent samples of those obtained to generate a simile of a GWAS which we could use to guide the feature selection process. We then took subsets of the remaining 100,000 individuals with increasing sizes, from 500 individuals up to 10,000 individuals. We then do 5-Fold Cross-Validation splitting those subsets into training and testing sets.

Furthermore, we took the simulation and extended it. Based on the assumption that we could be lacking more subjects to train in a better way the Machine Learning algorithms we decided to augment the existing data with artificial individuals obtained from a simulation with statistical characteristics similar to the ones found in the studies.First we reduced the number of SNPs to the amount we had present both in the IGAP results which included Odds Ratio as well as in the WGS. We then applied a clump as above without using any p-value filtering to obtain those SNPs in Linkage Equilibrium (As the simulator generates samples where all SNPs are in Linkage Equilibrium). The odds ratio associated to the disease were then obtained from the IGAP study, while the the allele frequencies were calculated from the full ADNI WGS (808 individuals, discarding the individuals related to each other). In this way we generated artificial data that is similar in terms of allele frequencies to the one present in the ADNI WGS as well as being similar with the results of the IGAP study with respect to the odds ratio of the disease.

Thus we generated with PLINK a set of sample using these metrics. Different sizes of data sets where used, having sets of 500, 5000 and 50000 individuals, this with the objective to validate the impact in the performance of the machine learning algorithms when using data sets with an increasing number of samples. The algorithms were trained exclusively on the artificial data, and were then tested with 138 individuals from the ADNI WGS data set which were not present in the IGAP study to ensure no information leakage was occurring, as well as the complete 471 individuals in the binary classification. The first subset is split 78% cases and 22 % controls, thus the ROC Score gives a much clearer view of the classification performance.

### 3.1 Validation Methodology

The main variable to analyze is the area under the Receiver-Operating-Characteristic, Area under the Curve (ROC-AUC) Score. 5-Fold Cross-Validation is used in the data set to ensure the results are statistically relevant and the validation does not over fit. This is done both in the direct case as well as in the simulation of a complex genetic disease. When testing the IGAP samples for train and the non-IGAP samples for test it is just done directly without Cross-Validation. Th For the case where we do the data-augmentation process we do the training on the different subsets and then test directly on the 138 individuals of the ADNI that are unrelated or in the whole subset of the 471 individuals.

## 4 Results

The first analysis done was using the ADNI data set directly with a varying number of significant SNPs. The SNPs obtained from the Clump file previously obtained were used as inputs, with a varying number of SNPs in order of significance being considered. The ADNI data set is split in test and training using 5-fold Cross Validation and the resulting values are shown in Figure 1, where it can be appreciated the maximum ROC AUC score value a machine learning method obtained was 0.66 when using around 20 SNPs using the Random Forest method.

**Fig. 1.**
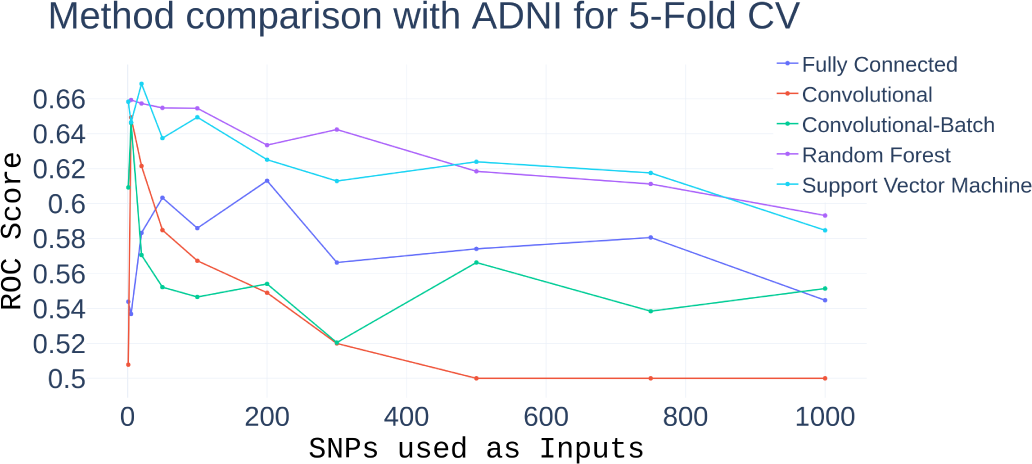
Method comparison using the Complete ADNI dataset. Analysis of the performance in terms of ROC AUC Score of the different classification methods when increasing the number of SNPs used as inputs, using 5-fold CV with the complete ADNI dataser. The SNPs used are in a descending order of statistical importance, with lower p-values as the first SNPs.

To further refine the results, a second experiment is made where we obtain the resulting ROC AUC score when selecting only those individuals who appear both in the ADNI data set and in the IGAP study as training set, while using the 138 individuals who were not included in the IGAP as a test set to ensure no information leakage occurs. We consider this as the Split analysis and the subsets as the IGAP-Independent case. In this case the resulting ROC AUC scores can be seen in Figure 2, where the results are more conservative than before. This could be due to the a-priori information, the class imbalance or the small sample size.

**Fig. 2.**
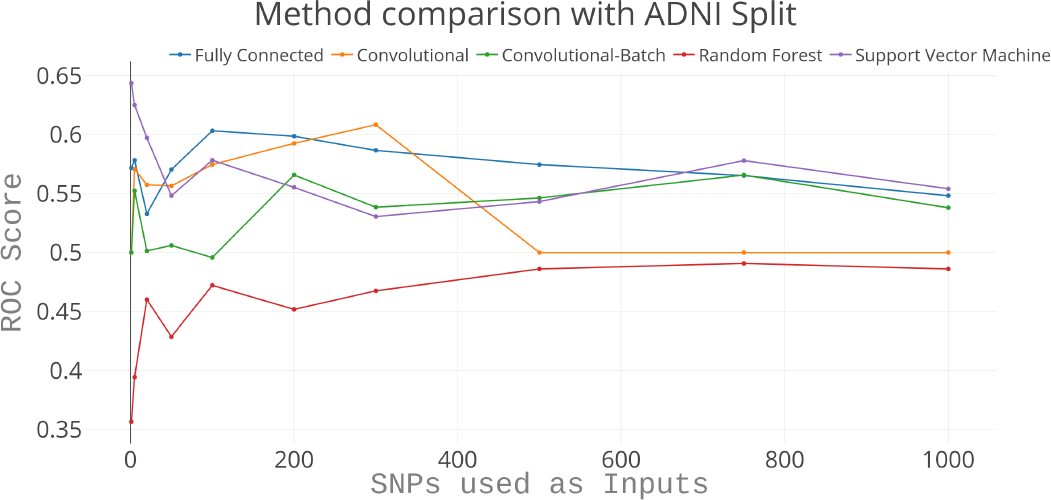
Method comparison using the Split ADNI dataset. Analysis of the performance in terms of ROC AUC Score of the different classification methods when increasing the number of SNPs used as inputs, training the model on the ADNI1 and ADNI GO samples and testing it in the IGAP-independent ADNI 2 Samples.

The next step taken was to attempt to simulate a disease and find out the performance of the methods when utilizing a larger data set. With the simulation we can clearly see that the use of a higher number of data samples leads to a much more precise classification as shown in Figure 3. And more interestingly, the increase in the number of samples and makes it so that using a higher number of SNPs as input becomes more valuable. Thus with small data sets it makes sense to use the highest-rated SNPs, but by introducing more SNPs in large data sets the results are refined further and a more precise classification can be achieved. This can be seen in Figures 4,5 and 6. For a direct contrast between the performance using two different subsets (500 and 10,000 respectively) Figure 7 and 8 can be analyzed to see how the performance increase is substantial.

**Fig. 3.**
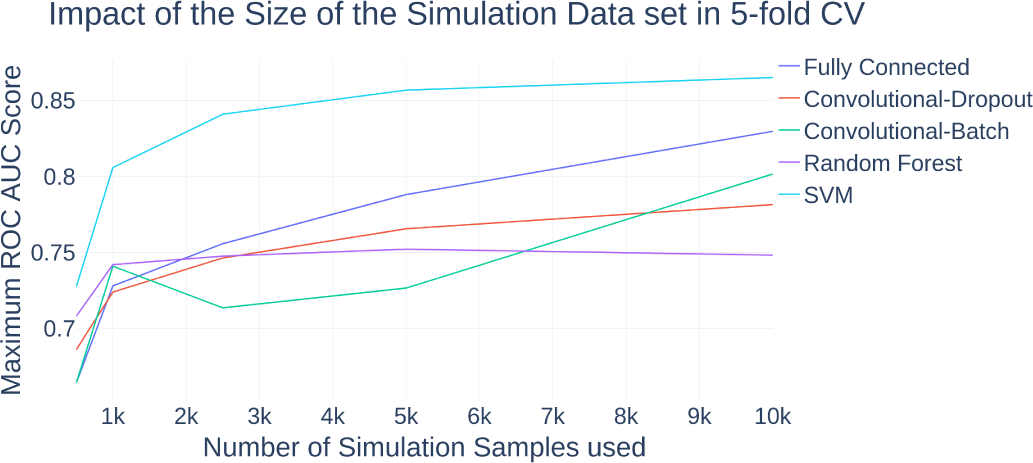
Impact of the subset size on the prediction. Analysis of the effect in the ROC AUC Score done by increasing the number of samples used for the 5-fold CV in the simulated dataset using the different classification methods.

**Fig. 4.**
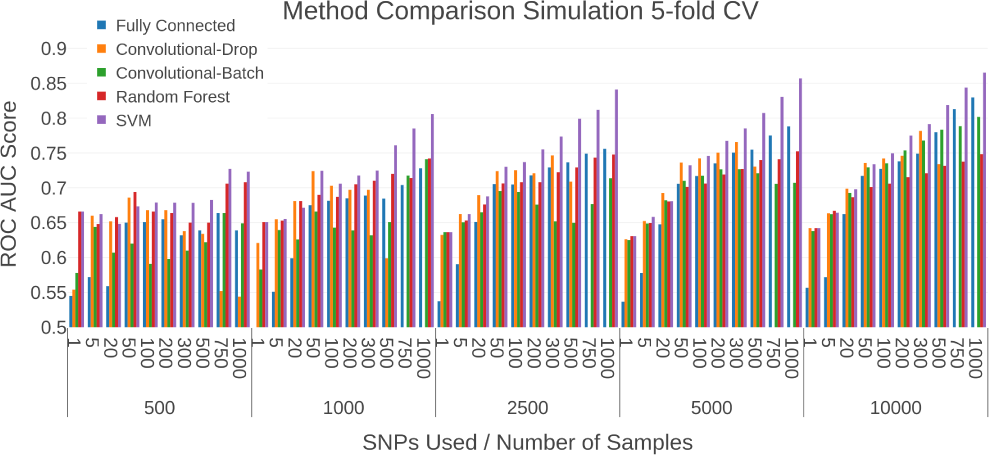
Method Comparison with Simulated Dataset. Comparison between the ROC AUC Score obtained using the different classification methods. The X axis firstly describes an increase of the number of SNPS used for classification, and afterwards an increase in the size of the simulation dataset

**Fig. 5.**
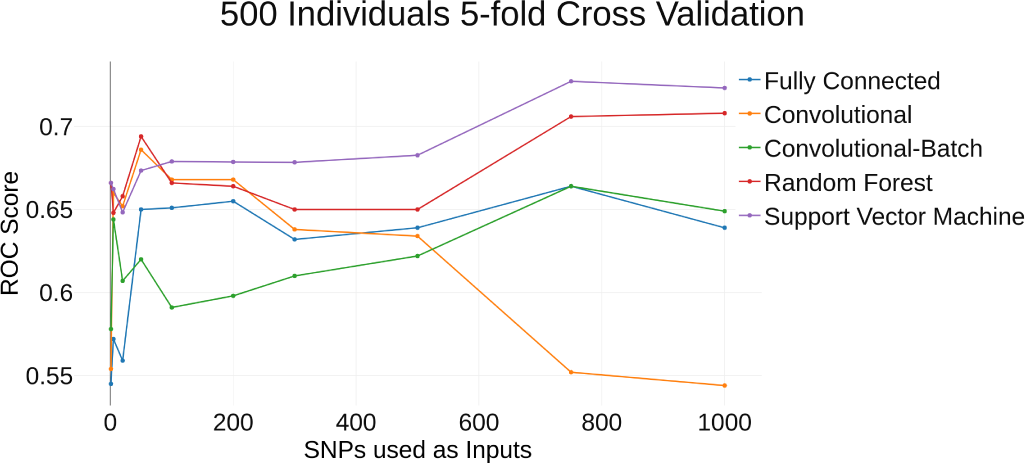
Method comparison with 500 Individuals. Comparison of the ROC AUC Score from 5-fold CV using different classification methods while increasing the number of SNPs given a simulation dataset with 500 Individuals

**Fig. 6.**
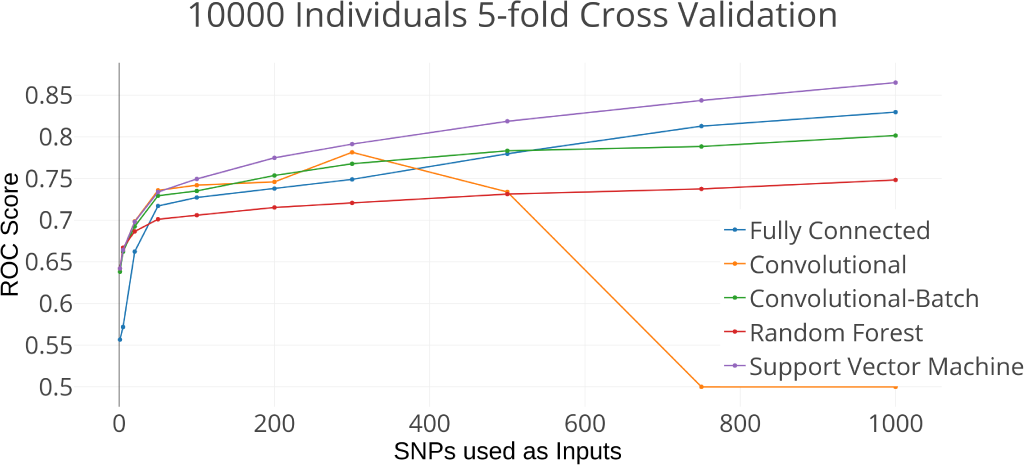
Method comparison with 10000 Individuals. Comparison of the ROC AUC Score from 5-fold CV using different classification methods while increasing the number of SNPs given a simulation dataset with 10,000 Individuals

**Fig. 7.**
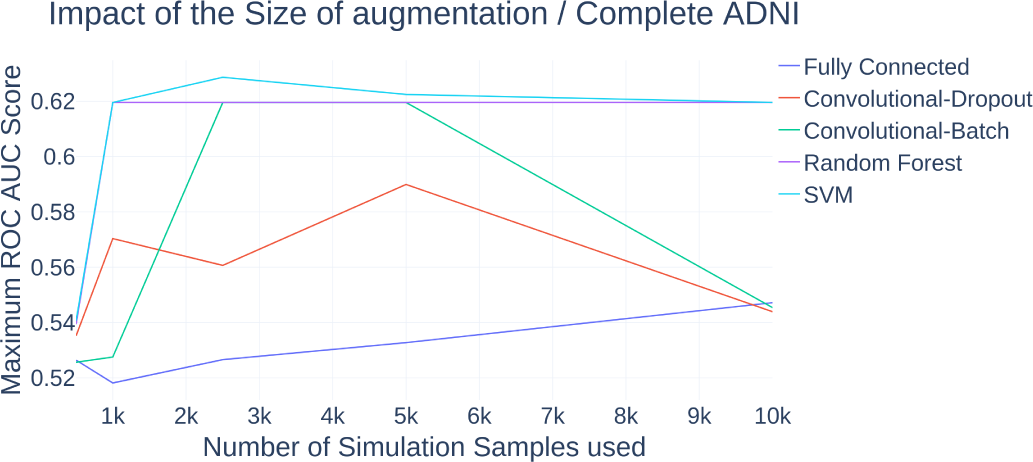
Impact of the augmentation size on the prediction. Analysis of the effect in the ROC AUC Score done by increasing the number of data-augmentation samples used for the training segment validated on the complete ADNI Dataset using the different classification methods.

**Fig. 8.**
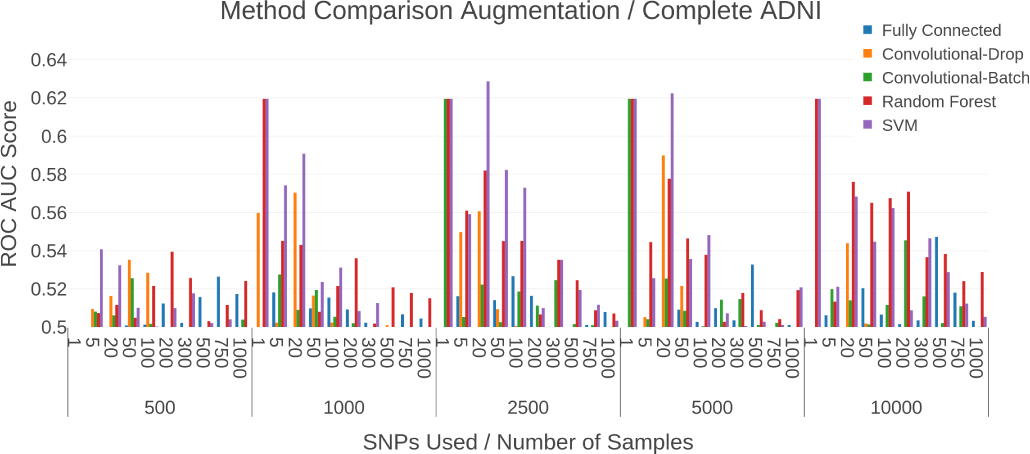
Method Comparison with ADNI Dataset. Comparison between the ROC AUC Score obtained using the different classification methods. The X axis firstly describes an increase of the number of SNPS used for classification, and afterwards an increase in the data-augmentation size validated on the complete ADNI dataset

The results from the previous simulation show that the use of more samples should benefit the prediction in the Alzheimer’s task. Thus we generated the artificially augmented data set for training purposes and used the ADNI data set as our test set. In this case the increase in the number of samples did not directly mean a better performance and a higher use of SNPs, as we can see once we start using over 100 SNPs the performance on the ADNI subset tends to decay. Plus, the increase in size of the training data set also did not mean constant increases as in the previous scenario. We can see that the use of more SNPs coupled with a decent size of artificially-augmented samples gives the best results on the ROC Score, which shows that both of these factors have a role to play in the prediction. The convolutional dropout network did have some issues when using too much SNPs and dropped to a classification, and in general the simpler models such as Random Forest and Support Vector Machine performed better. This shows that there is only so much that can be accomplished with data augmentation, and that the disease might actually be focused on a smaller number of uncorrelated genes as supposed in the simulation. Figures 7 and 8 illustrate these results.

Further restrictions were imposed on the data set used for testing, as some of the ADNI samples were also taken into account for the IGAP study. Thus we restricted the testing to those individuals who did not participate in the IGAP study. For this case we can see in Figures 9 and 10 that due to the small sample size and the class imbalance the results are lower as the previous data set. Specifically they tend to show good results using few amounts of SNPs (APOE4 mainly) and the increase in sample size used to train does not represent an increase in the prediction capability. This gives us the impression that the learning models either do not generalize correctly, or the few individuals from the unbalanced data set do not correspond to the genetic markers learned from the IGAP-based simulation. Comparing these results with the ones obtained from the ADNI split the results are in average better for the data augmentation process than from the standard.

**Fig. 9.**
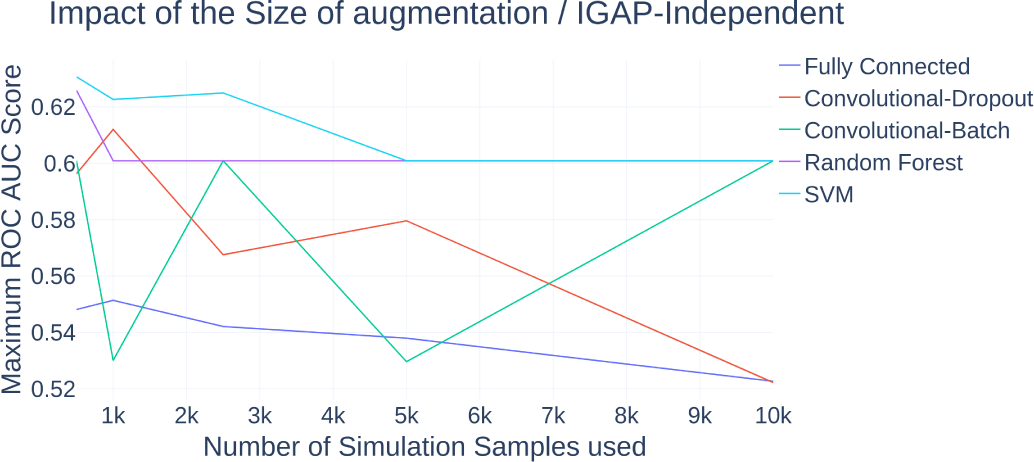
Impact of the augmentation size in IGAP-Independent Subset. Analysis of the effect in the ROC AUC Score done by increasing the number of data-augmentation samples used for the training segment validated on the IGAP-Independent subset using the different classification methods.

**Fig. 10.**
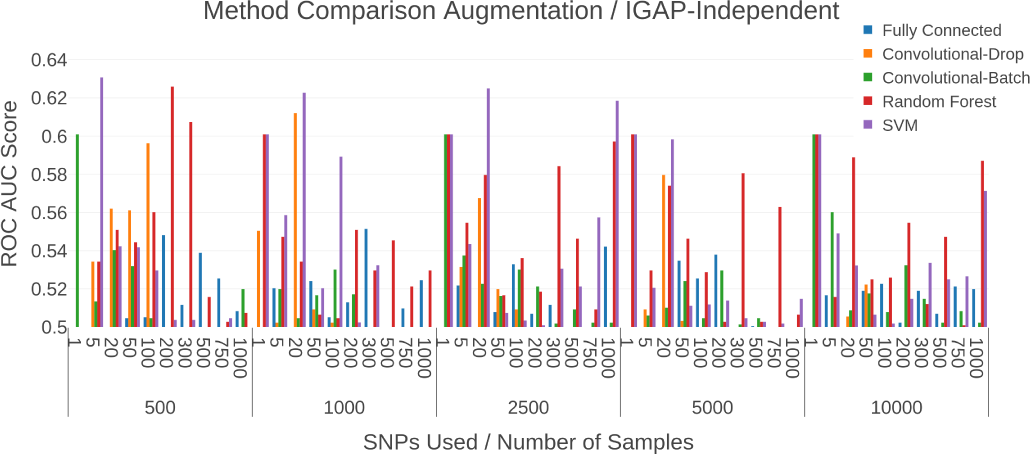
Method Comparison with IGAP-Independent Subset. Comparison between the ROC AUC Score obtained using the different classification methods. The X axis firstly describes an increase of the number of SNPS used for classification, and afterwards an increase in the data-augmentation size validated on the IGAP-Independent subset.

## 5 Discussion

Previous research on the early detection of late-onset Alzheimer’s disease have relied on a variety of clinical biomarkers for disease prediction [2]. The efficacy of experimental treatments and palliative interventions rely heavily on the early detection of the disease [5]. Unfortunately, clinical biomarkers such as beta amyloid and tau proteins are correlated with disease progression. Therefore, their usefulness for the early detection of the disease remains controversial.

The etiology of LOAD is likely to be motivated by both environmental and genetic components. However, the genetic component seems to a major determinant as the heritability of the disease has been estimated to be ∼ ∼80% [21]. Therefore, genetic testing hold the potential to provided sufficiently accurate predictions of the disease using genetic data exclusively. Unfortunately, the genetic variants with associations with LOAD discovered by GWAS studies are only capable to explain a fraction of this genetic component (33%). Therefore, methodologies that account for this missing heritability are required to achieve better predictions [9] [8].

In this work, we propose the use of deep neural networks with dense layers for predicting late-onset Alzheimer’s disease from genetic data. Our work hypothesis postulated that the used of a large number of genetic variants would allow us to improve the classification performance of the proposed model. We expect the deep learning model to create hierarchical features with the potential to account for the missing heritability of the disease.

Experimental results indicate that classification performance of ∼65% AUC can be achieved with the proposed model. In comparison, the use of the APOE4 gene with our dataset gives a predictive score of 0.61% 0.65% on the Cross-Validation and 56% 64% on the Split Validation depending on the method. Most importantly, the experiments reported here suggest that an increasing number of genetic variants as predictors hold the potential to contribute to improve the predictive capabilities of the proposed model providing that a sufficiently large number of samples are available.

In the majority of experiments reported here, random forest produced better results that deep learning models. However, according to empirical observations on the performance of deep learning models in our experiments, we expect the latter to outperform the former as more data becomes available. In general, deep learning models have shown to scale the performance better than other machine learning models with increasing amounts of data [13].

LOAD prediction is a challenging problem. In effect, the etiology and the genetic architecture of the disease remain unexplained. Moreover, accurate diagnosis of the disease is still an open problem. Therefore, labeled datasets including confirmatory diagnosis are not currently available. This data would be critical for the construction of accurate predictive models.

In addition, it is unclear whether or not there are useful genetic variants that account for the transition from mild cognitive impairment to LOAD. According to recent studies, currently available LOAD polygenic scores are not capable to predict mild cognitive impairment to LOAD progression [14]. Therefore, alternative models are also required for the accurate prediction of disease progression.

## 6 Conclusion

This study proposes the use of deep neural networks for the prediction of late-onset Alzheimer’s disease from a large collection of genetic variants. Experimental results indicate that the proposed model holds promise to produce useful predictions for clinical diagnosis of LOAD.

## Acknowledgments

We thank our colleagues from the Bioinformatics for Clinical Diagnosis Research Program, School of Medicine and Health Sciences, Tecnologico de Monterrey, for their valuable comments on this work. This work was supported by Tecnologico de Monterrey.

Data collection and sharing for this project was funded by the Alzheimer’s Disease Neuroimaging Initiative(ADNI) (National Institutes of Health Grant U01 AG024904) and DOD ADNI (Department of Defense award number W81XWH-12-2-0012). ADNI is funded by the National Institute on Aging, the National Institute of Biomedical Imaging and Bioengineering, and through generous contributions from the following: AbbVie, Alzheimers Association; Alzheimers Drug Discovery Foundation; Araclon Biotech; BioClinica, Inc.; Biogen; Bristol-Myers Squibb Company; CereSpir, Inc.; Cogstate; Eisai Inc.; Elan Pharmaceuticals, Inc.; Eli Lilly and Company; EuroImmun; F. Hoffmann-La Roche Ltd and its affiliated company Genentech, Inc.; Fujirebio; GE Healthcare; IXICO Ltd.; Janssen Alzheimer Immunotherapy Research & Development, LLC.; Johnson & Johnson Pharmaceutical Research & Development LLC.; Lumosity; Lundbeck; Merck & Co., Inc.; Meso Scale Diagnostics, LLC.; NeuroRx Research; Neurotrack Technologies; Novartis Pharmaceuticals Corporation; Pfizer Inc.; Piramal Imaging; Servier; Takeda Pharmaceutical Company; and Transition Therapeutics. The Canadian Institutes of Health Research is providing funds to support ADNI clinical sites in Canada. Private sector contributions are facilitated by the Foundation for the National Institutes of Health (www.fnih.org). The grantee organization is the Northern California Institute for Research and Education, and the study is coordinated by the Alzheimers Therapeutic Research Institute at the University of Southern California. ADNI data are disseminated by the Laboratory for Neuro Imaging at the University of Southern California.

We thank the International Genomics of Alzheimer’s Project (IGAP) for providing summary results data for these analyses. The investigators within IGAP contributed to the design and implementation of IGAP and/or provided data but did not participate in analysis or writing of this report. IGAP was made possible by the generous participation of the control subjects, the patients, and their families. The iSelect chips was funded by the French National Foundation on Alzheimer’s disease and related disorders. EADI was supported by the LABEX (laboratory of excellence program investment for the future) DISTALZ grant, Inserm, Institut Pasteur de Lille, Universit de Lille 2 and the Lille University Hospital. GERAD was supported by the Medical Research Council (Grant n 503480), Alzheimer’s Research UK (Grant n 503176), the Wellcome Trust (Grant n 082604/2/07/Z) and German Federal Ministry of Education and Research (BMBF): Competence Network Dementia (CND) grant n 01GI0102, 01GI0711, 01GI0420. CHARGE was partly supported by the NIH/NIA grant R01 AG033193 and the NIA AG081220 and AGES contract N01AG12100, the NHLBI grant R01 HL105756, the Icelandic Heart Association, and the Erasmus Medical Center and Erasmus University. ADGC was supported by the NIH/NIA grants: U01 AG032984, U24 AG021886, U01 AG016976, and the Alzheimer’s Association grant ADGC10196728.

